# Genetic Insights into Agronomic and Morphological Traits of Drug-Type Cannabis Revealed by Genome-Wide Association Studies

**DOI:** 10.1101/2023.11.09.566286

**Authors:** Maxime de Ronne, Éliana Lapierre, Davoud Torkamaneh

## Abstract

Cannabis sativa L., previously concealed by prohibition, is now a versatile and promising plant, thanks to recent legalization, opening doors for medical research and industry growth. However, years of prohibition have left the cannabis research community underdeveloped and lacking knowledge about cannabis genetics and trait inheritance. To bridge this gap, we conducted a comprehensive genome-wide association study (GWAS), using a panel of 176 drug-type cannabis accessions, curated to represent the Canadian legal market. This pioneering GWAS harnessed the power of high-density genotyping-by-sequencing (HD-GBS), resulting in an exhaustive catalog of 800K genetic variants. These variants served as the bedrock for a GWAS designed to dissect the genetic foundations of nine key traits. To identify the most robust markers associated with these traits, two sophisticated statistical methodologies were used (SUPER and BLINK), ultimately identifying 33 markers significantly associated with agronomic and morphological traits. Several identified markers exert a substantial phenotypic impact, guided us to a rich trove of putative candidate genes that reside in high linkage-disequilibrium (LD) with the markers. These markers show great promise for revolutionizing cannabis breeding to meet diverse needs. In doing so, they lay the solid foundation for an innovative cannabis industry poised to reshape the future.

## Introduction

Cannabis (*Cannabis sativa* L.), an annual and dioecious plant species belonging to the Cannabaceae family, stands as one of the earliest domesticated plants. Its rich history is intertwined with the socioeconomic and cultural development of human societies^1,2^. This versatile crop has served a multitude of purposes, offering valuable fibers for ropes and nets, abundant production of protein- and oil-rich seeds, applications in traditional medicine dating back to approximately 8,000 BCE, and psychoactive properties^3^. In Canada, the trajectory of cannabis cultivation took a significant turn, transitioning from a 1920s prohibition to the legalization of hemp cultivation in 1998, followed by the authorization of medical use in 2001 and recreational use in 2018^4,5^. Despite the fact that cannabis is known to produce over 545 potentially bioactive secondary metabolites, in Canada, the USA and the Europe, it is legally categorised based on the concentration of a single cannabinoid, the Δ^9^-tetrahydrocannabinol (THC), present in the female flowers^6^. Cannabis plants with less than 0.3% THC are classified as hemp-type, while those with greater than 0.3% THC are labeled as drug-type cannabis. The shift in legislation has fueled the development of diverse industries, significantly contributing to Canada’s gross domestic product (GDP) and job market, injecting approximately $43.5 billion into the economy and creating over 151,000 jobs in four years (2018-2022)^7^. The historical and societal significance of cannabis is undeniable, and recent changes in legislation worldwide have propelled it into the forefront of scientific investigation, research and development^8^. Since the discovery of THC in 1964, extensive efforts have been made to characterize the metabolome of hundreds of cannabis plants, leading to discovery of over 150 terpenoids, 120 cannabinoids and various flavonoids^9,10^. Likewise, there have been substantial strides in unraveling the cannabis genome and creating a worldwide *C. sativa* genomics resource^11,12^. Notably, significant progress in cannabis genome assembly has been achieved through the utilization of long-read sequencing technologies (i.e., PacBio and Oxford Nanopore Technologies) coupled with scaffold anchoring with genetic linkage maps and the integration of Hi-C data. These advancements have led to the development of four chromosome-level assemblies^13,14^. Among them, the cs10 v2 assembly (GenBank acc. no. *GCA_900626175.2*) is considered as the most complete and has been proposed as the reference genome for cannabis by the International Cannabis Research Consortium (ICGRC)^15^. In this assembly, the *C. sativa* has been estimated to be around 875.7 Mb, characterized by a pair of sex chromosomes and nine autosomes, comprising 31,170 annotated genes^16^. The *de novo* assembly of cannabis genomes was fraught with challenges due to a substantial level of heterozygosity (ranging from approximately 12.5 to 40.5%), and a remarkable abundance of repetitive elements, accounting for roughly 70% of the genome^3^. The in-depth characterization of the metabolome and genome of *C. sativa* provided new opportunities for medical research, industrial growth and the development of modern agronomic practices.

Despite these progresses, the 20^th^ century prohibition of cannabis has hindered its cultivation from fully benefiting from the tools introduced during the Green Revolution^5^. For many years, cannabis breeding occurred in clandestine operations, relying on undocumented methods and a dearth of modern technologies. Similar to other high-value crops, modern breeding technologies hold the promise of enhancing cannabis traits to meet diverse needs, spanning manufacturing, medicinal, recreational, and culinary uses ^17^. Cannabis research community is hugely undersized and suffers from a scarcity of understanding of cannabis genetics and how key traits are expressed or inherited^18^. Thus, a better understanding of the genetic basis of agronomic and morphological traits of drug-type cannabis appears to be a prerequisite for the development of improved cannabis varieties, optimizing cultivation practices, and conserving valuable genetic resources^3^.

The advent of next-generation sequencing technology (NGS)^19^, which offers cost-effective high-throughput sequencing, coupled with the availability of powerful bioinformatic tools^20,21^, have facilitated the widespread adoption of genotype-phenotype association studies to investigate the relationship between genetic variation and phenotypic traits for a wide range of crops^22^. Recent classic quantitative trait loci (QTL) mapping studies have enabled identification of maturity-related QTL in both hemp^23^ and drug-type cannabis^24^. Classic QTL mapping analysis defines molecular markers linked to a phenotype segregating within parental lines, in contrast to modern genome-wide association studies (GWAS) which identify loci related to phenotypes within large populations of unrelated individuals^22^. GWAS use the information of linkage disequilibrium (LD) between a QTL and neighboring genetic markers to identify the regions on the genome that influence traits. However, when applied to a large set of individuals, the sequencing cost remains the most limiting factor, especially in heterozygous organisms like cannabis where a high sequencing depth per sample is needed to accurately determine genotypes^25^. To address this challenge, cost-effective high-throughput genotyping methods (e.g., restriction-site associated DNA sequencing (RAD-Seq)^26^, genotyping-by-sequencing (GBS)^27^ and High-Density GBS (HD-GBS)^28^, based on reduced-representation sequencing approaches (RRS; Poland and Rife, 2012), have been developed. Recent GWAS studies in hemp-type cannabis^29–31^ to investigate fiber quality, flowering time and sex determination and drug-type cannabis^32^ to investigate genetic basis of terpenes have enabled identification of significant genetic markers. The newly identified QTL will enable the early selection of promising individuals through marker-assisted selection (MAS)^33^, thereby reducing the labor and costs associated with development of improved varieties. Genetic association studies are, therefore, of significant value in advancing breeding programs towards molecular approaches^22^.

While flowering time and sex determination have been focal points in cannabis breeding, the genetic basis of other important agronomic traits (e.g., yield, height, days to maturity, etc.) remain largely unexplored. Morphological traits should be duly considered due to their established intercorrelations with yield, maturity and cannabinoid profiles^34^. For instance, cannabis plants cultivated for medicinal and recreational application exhibit shorter stature, have thinner stems, more nodes, higher floral density, and a different cannabinoid profiles compared to industrial hemp plants^35^. On the other hand, genetic backgrounds that prioritize yield may negatively impact THC production, and vice versa^34^. Investigating genetic variations associated with agronomic and morphological traits is essential for establishing the genetic groundwork for developing tailor-made cannabis varieties, along with breeding tools such as MAS and genomic selection (GS)^36^.

To facilitate the development of molecular tools for cannabis breeders and researchers, the present study provides high-value markers linked to essential agronomic and morphological traits, identified through GWAS conducted on 176 drug-type cannabis accessions representative of the Canadian legal market. High-confidence markers associated with essential traits were cross-validated using robust single- and multi-locus statistical methods. In summary, this study lays the groundwork for a comprehensive understanding of the genetic foundations underpinning the agronomic and morphological traits in cannabis. The markers identified through this research promise to significantly expedite breeding efforts, empowering us to cultivate cannabis varieties optimized for various purposes and applications.

## Experimental procedures

### Plant material and Phenotyping data

All research activities, including the procurement and cultivation of cannabis plants, were executed in accordance with our cannabis research license (LIC-QX0ZJC7SIP-2021) and in full compliance with Health Canada’s regulations. In total, in this study, we used 176 drug-type accessions each accompanied by phenotyping data sourced from Lapierre et al. (2023b). These accessions were thoughtfully selected from diverse sources to ensure representation of the broad spectrum of the drug-type cannabis varieties available in the legal market of Canada (Supplementary Table 1).

In this study, we used four key productivity-related traits, including fresh biomass (FB), dried flower weight (DFW; representing yield), sexual maturity (SM) and harvest maturity (HM; days to maturity). Additionally, we included five morphological traits, namely stem diameter (SD), canopy diameter (CD), height, internode length index (ILI) and node counts (NC). It is worth noting that values were originally recorded in inches were converted to centimeter for consistency. Histograms representing the distribution of each trait for the 176 accessions were generated using R v4.2.1^37^ with the ‘*hist*’ function. Furthermore, a *t*-test was performed to determine whether the minimum and maximum values of each trait significantly differed from the overall population mean.

### Sequencing and genotyping

#### DNA isolation, library preparation and sequencing

Approximately 50 mg of young leaf tissue from each accession was collected for DNA extraction. The collected leaf tissues were air-dried for four days using a desiccating agent (Drierite; Xenia, OH, USA) and then ground with metallic beads in a RETSCH MM 400 mixer mill (Fisher Scientific, MA, USA). DNA extraction was carried out using the CTAB-chloroform protocol. In brief, the powdered tissue was treated with a CTAB buffer solution, followed by a phenol-chloroform extraction procedure. The resulting DNA pellet underwent ethanol washing and was subsequently re-suspended in water. DNA quantification was carried out using a Qubit fluorometer with the dsDNA HS assay kit (Thermo Fisher Scientific, MA, USA), and concentrations were adjusted to 10 ng/μl for all samples. Final DNA samples were used to prepare HD-GBS libraries with *Bfa*I as described in Torkamaneh *et al.* (2021) at the Institut de biologie intégrative et des systèmes (IBIS), Université Laval, QC, Canada. Sequencing was conducted on an Illumina NovaSeq 6000 (Illumina, CA, USA) with 150 paired-end reads at the Genome Quebec Service and Expertise Center (CESGQ), Montreal, QC, Canada.

#### SNP calling and filtration

Sequencing data were processed with the Fast-GBS v2.0^38^ using the *C. sativa* cs10 v2 reference genome (GenBank acc. no. *GCA_900626175.2*)^13^. For variant calling a prerequisite of a minimum of 6 reads to call a single nucleotide polymorphism (SNP) was opted. Raw SNP data were filtered with VCFtools^39^ to remove low-quality SNPs (QUAL <10 and MQ <30) and variants with proportion of missing data exceeding 80%. Missing data imputation was performed with BEAGLE 4.1^40^, followed by a second round of filtration, retaining only biallelic variants with heterozygosity less than 50% and a minor allele frequency (MAF) of > 0.06. Additionally, variants residing on unassembled scaffolds were removed. The resulting catalog of ∼282K SNPs was used to conducted genetic analysis, population structure assessment and GWAS.

### Genetic analysis

#### Marker description

Read counts and coverage were calculated with SAMtools “coverage” parameter^41^. Proportion of heterozygous variants, MAF and Tajima’s D value were estimated using TASSEL5^42^. The proportion of SNPs located within annotated genes was determined with BEDTools^43^ by analyzing the number of SNPs overlapping with gene regions^44^. To visualize the distribution of SNP density, a plot was produced with rMVP^21^ using ‘*plot.type=”d”*’ parameter, in combination with the gene density distribution.

#### LD decay and Haplotype block

Pairwise-LD was calculated with PLINK v1.9^45^ using ‘*--r2 --ld-window-r2 0*’ parameters. LD, measured as the allele frequency correlation (r^2^), was determined for all pairwise SNPs within each chromosome. The LD decay was determined by plotting the r^2^ values against the distance of SNP pairs, using an R script adapted from Remington *et al.* (2001). The point of intersection between the LD curve and the predefined r^2^ threshold determined the LD decay. Estimation of haplotype blocks (HBs) was performed with PLINK v1.9 using ‘*-- blocks no-pheno-req --ld-window-kb 999*’. A *t*-test was conducted in R to assess whether the LD decay of the chromosome X significantly differed from that of other chromosomes.

### Population structure analysis

#### Population structure and admixture

Population admixture was determined using a variational Bayesian inference algorithm implemented in FastStructure^47^ for a number of subpopulations (*k*) set from 1 to 10. The optimal number of *k* explaining the population complexity was estimated using the ChooseK tool from FastStructure and admixture proportions were visualized using Distruct v2.3. The kinship matrix (K*) was generated using TASSEL5 with the Centered_IBS method and plotted with GAPIT v3^20^.

#### Discriminant analyses of principal components (DAPC) for population structure

Population structure was further investigated using discriminant analyses of principal components (DAPC)^48^ using the R package ‘*adegenet*’. The number of cluster was estimated using ‘*find.cluster*’ function with a maximum limit set to 40 clusters and 200 principal components (PCs). Optimal number of clusters was determined by the minimal Bayesian Information Criterion (BIC) value for different number of *k* (i.e., *k*=3, Supplemental Figure S1ab). To visualize the DAPC using the ‘*scatter*’ function, the optimal number of PCs was estimated with two cross-validation procedures using ‘*optim.a.score*’ (i.e., PCs=20, Supplemental Figure S1c) and ‘*xvalDapc*’ (i.e., PCs≤20, Supplemental Figure S1d).

#### Comparison of population assignments and trait analysis

Cluster assignments from both FastStructure and DAPC were compared using the ‘*table*’ function for a *k* value of 3 and 6. An analysis of variance (ANOVA) and permutational ANOVA (PERMANOVA) were performed for traits following and deviating from the normal distribution, respectively, using the ‘*adonis2*’ function from R package ‘*vegan*’. Cluster assignments obtained from FastStructure were used as covariate. In cases where ANOVA/PERMANOVA indicated a significant difference, the post-hoc Tukey honestly significant difference (HSD) test was performed to determine which pairs were significantly different. Violin plots were generated with ‘*ggplot2*’ in R and Tukey significant differences were represented by letter.

### Genome-wide association analysis

Marker-trait association analysis was performed using GAPIT v3^20^, using the 282K high-quality SNPs and the phenotyping data for nine different traits. To ensure the identification of high-confidence SNPs while minimizing false positives, a cross-confirmation of two complementary methods, Settlement of MLM Under Progressively Exclusive Relationship (SUPER)^49^ for single-locus analysis and Bayesian-information and linkage-disequilibrium iteratively nested keyway (BLINK)^50^ for multi-locus analysis, were used^51,52^. Both methods incorporated population structure (i.e., P matrix generated with FastStructure for *k*=3) and kinship (i.e., K* matrix generated with TASSEL5) for the analysis. The threshold of significance for marker-trait associations in both methods was set to ensure a false discovery rate < 0.05, adjusted with a Benjamini-Hochberg correction. However, when a significant marker was commonly identified by both methods, the BLINK results were reported. Markers with a phenotypic variance explained (PVE) less than 3% were excluded from the analysis as they were considered uninformative and of limited interest to breeders. Manhattan plots showing –log10(*p*) distribution of markers by chromosome were generated with rMVP^21^ using ‘*plot.type=”m”*’ and quantile-quantile (QQ) plots were created with GAPIT v3^20^.

### Candidate gene identification

Due to the substantial genetic diversity present in cannabis, only a limited number of SNPs exhibited a strong LD (r²≥0.95). Therefore, to pinpoint genetic regions of interest, only markers in high LD (r²≥0.75) with significant markers were retained to define haplotype blocks (HBs). Markers failing to form HB and residing outside of genetic regions were removed from the candidate gene investigation. Genes located in the HBs (defined by the 5’-most and 3’-most marker of the HB) were considered as candidate genes. The gene ontology (GO) annotations of these candidate genes were examined based on the description provided by the NCBI *Cannabis sativa* Annotation Release 100. To further confirm and provide a more detailed functional annotation of candidate genes, phylogenetic ortholog inferences were performed using OrthoFinder^53^ with the *Arabidopsis thaliana* transcriptome (TAIR 11)^54^.

## Results & Discussion

### A broad range of phenotypic variation among the 176 drug-type accessions

The population displayed significant phenotypic diversity (*p*<0.001) across the nine examined traits (Figure 1, Supplemental Table S1). For instance, FB exhibited a substantial variation, ranging from 90 to 1,260 g, while plant height varied between 22 to 109 cm. SM also showed a significant diversity, with individuals maturing between 20 to 68 days. With the exception of SM, all other traits displayed a unimodal distribution, suggesting a complex genetic control involving multiple QTL. Furthermore, these traits exhibited highly skewed distributions, indicating that some accessions may carry specific alleles or combinations of alleles exerting a substantial impact on these traits. This phenotypic diversity within the cannabis accessions provides a robust foundation for GWAS, aligning with established criteria for successful GWAS outcomes^22^.

**Figure 1.**
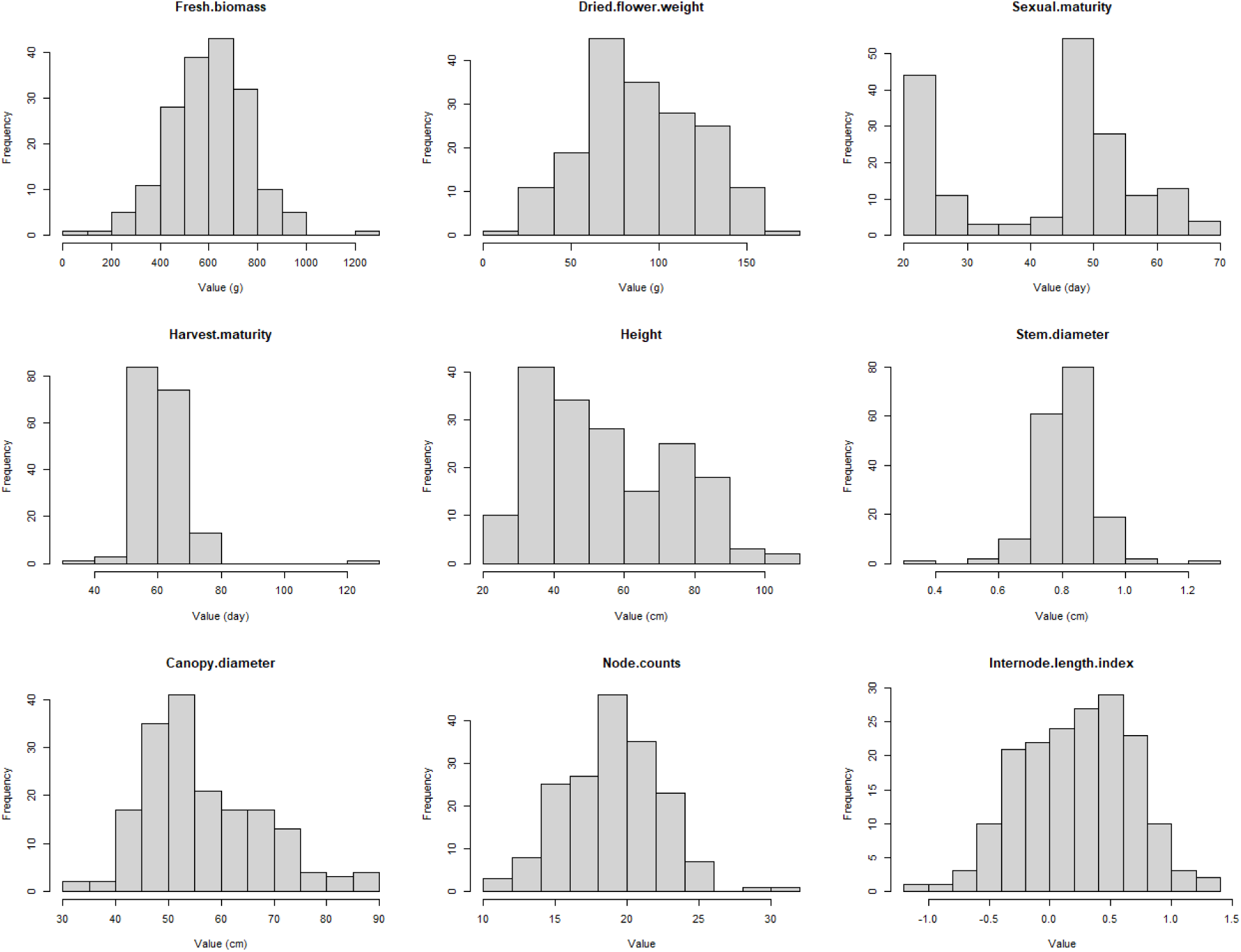
Frequency distribution of phenotypic data for 176 drug-type accessions used in this study.

### Genetic diversity in the GWAS-panel revealed by dense genotyping

To achieve comprehensive marker coverage across the cannabis genome, an HD-GBS approach was used. Sequencing of HD-GBS libraries generated 486M reads, averaging 2.8M reads per sample. This extensive sequencing effort resulted in an average per-sample coverage of 7.7% of the cs10 v2 assembly, achieving a remarkable cumulative coverage of 34.1% across the entire genome for the entire population. The analysis of variant calling from our sequencing data initially yielded a substantial dataset of 2.7M raw variants that met the quality criteria. Following filtering for missing data and minor allele frequency (MAF of 1%), we successfully identified ∼800K polymorphic variants. Subsequently, we performed a secondary round of filtering, primarily aimed at retaining common variants, as defined by a MAF of 6%, retaining approximately 39% of the raw data. While this filtering step may exclude rare variants that could potentially influence complex traits, it is essential to reduce the risk of false-positive associations and ensure that a minimum of 10 accessions carries the significant allele, thereby preventing overfitting in GWAS models^22^. The HD-GBS approach and the filtering procedures resulted in a catalog of 282K high-quality SNPs (all details in Supplemental Table S2). Within this catalog, 25.5% of the genotypes were found to be heterozygous and the SNPs exhibited an average MAF of 21.7%. For a detailed overview of filtering steps and the number of variants retained at each stage, refer to Supplementary Table S3. Overall, this SNP catalog represents an exhaustive genetic resource for the subsequent GWAS and underscores the robustness of the genotyping strategy used in this study.

Markers were exceptionally well distributed across the genome, ensuring coverage of gene-rich regions. On average, there was one marker per every ∼3 kb of the genome, which significantly enhances the likelihood of identifying markers in strong LD with putative candidate genes or regions (Figure 2a). Across the entire physical map, only 12 gaps exceeding 1 Mb, with the largest being 1.2 Mb, were identified. Comparing our dataset with the RAD-Seq method used in the study of Petit *et al.* (2020ab), by employing comparable filtration criteria, the HD-GBS approach yielded a comparable number of markers while utilizing only one-tenth of the sequencing efforts (averaging 2.8M vs. 29.7M reads per sample). Therefore, the density and genomic distribution of SNPs provided by the HD-GBS approach make it a cost-effective option for conducting GWAS on large cannabis panel. Furthermore, this approach is compatible with the miniaturization of sequencing libraries using the NanoGBS procedure, which further contributes to substantial cost reduction in genotyping^55^.

**Figure 2.**
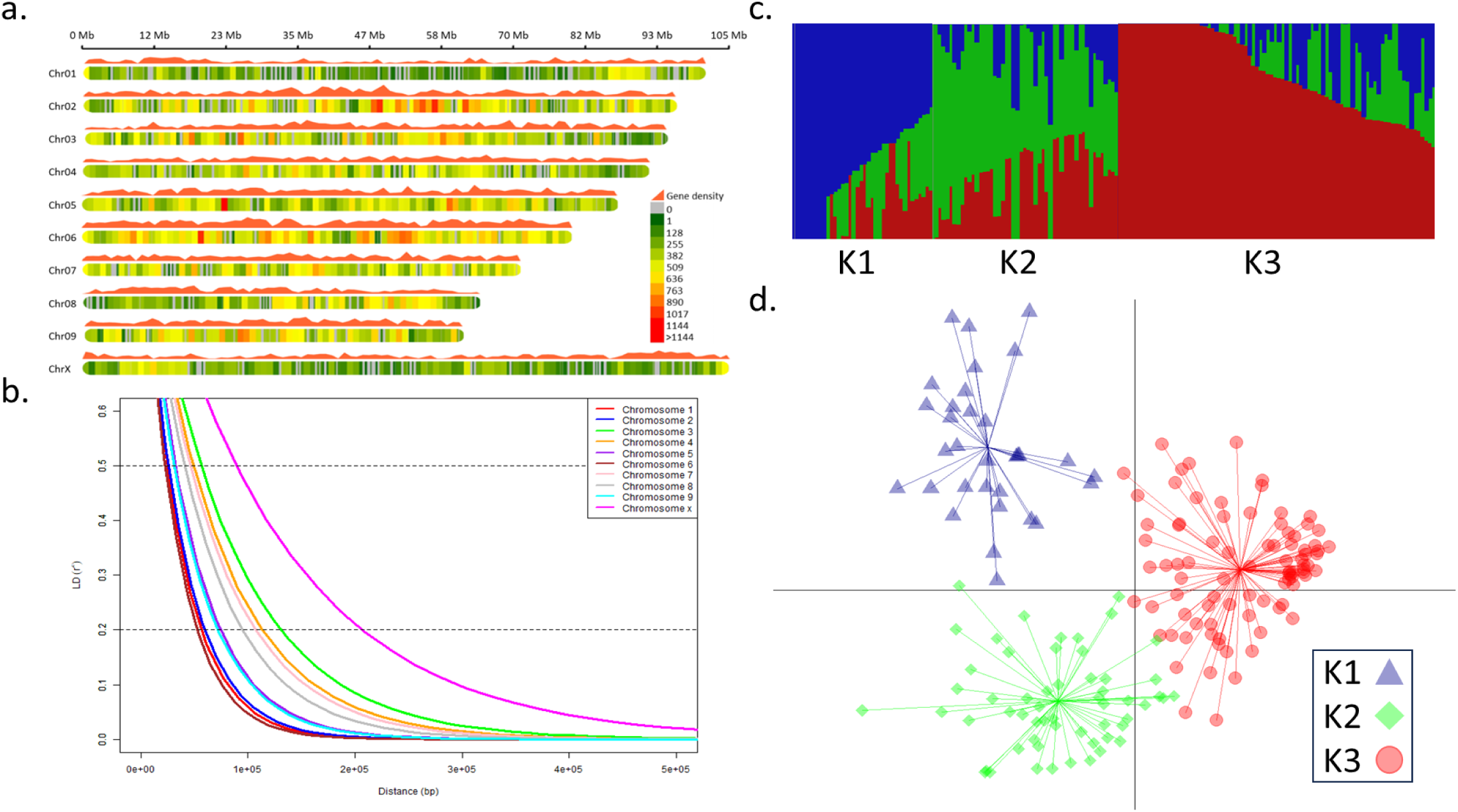
Genome-wide distribution of markers, linkage disequilibrium (LD) and population structure analysis. **a.** Density plot of markers and genes across the genome. Colors represent the number of SNPs within 1 Mb window size. **b.** LD decay in each chromosome where LD values of intra-chromosomal pairwise markers were ploted against physical distance. **c.** Admixture plot for k=3 using FastStructure. The vertical lines represent the accessions, and the y-axis represents the probability that an individual belongs to a subgroup. **d.** Discriminant analysis of principal components (DAPC) scatter plot showing population structure.

The average extent of LD decay to its half ranged from 22.6 to 89.0 kb across different chromosomes (Figure 2b). It is important to note that LD decay is a relative value and does not precisely reflect to reality recombination rates throughout the entire genome, particularly between hetero chromatic and euchromatic regions^56^. However, this measure proved valuable for comparing the impact of domestication and selection on recombination rates among the different populations. In this context, the LD observed in the GWAS-panel showed rapid decay compared to modern cultivars of comparable genome size, such as soybean (where LD may extend over 100 kb^57^) and tomato (where LD can extend over 1 Mb^58^). Nevertheless, LD decayed to its half more slowly compared to a recent study of 110 domesticated and landrace cannabis accessions from various worldwide origins, where LD decayed over approximately 10 kb^59^. This resulted in a large number of small HBs with an average size of ∼4 kb (Supplemental Table S3). It is worth noting that the LD decay on the sex chromosome was almost twice as high (*p*<0.001) compared to autosomes. These observations were consistent with the recent history of cannabis cultivation in Canada, characterized by extensive hybridization efforts by breeders with a particular focus on sexual characteristics, such as the production of female flowers^2^.

### Low level of population structure

The population structure within the GWAS-panel was assessed using the 282K high-quality SNPs. Initially, the degree of admixture of individuals and clustering inference was estimated by FastStructure. While the model maximizing the marginal likelihood suggested a *k* value of 6, the optimal number of principal components (PCs) to explain the structure of the population was determined to be 3. The *k* value of 3 revealed two sub-populations (sub-population 1 and 3, Figure 2c) with low admixture compared to a *k* value of 6 (Supplemental Figure S2), indicating a more robust assignments with more homogeneous individuals within each cluster. Using the BIC criterion, DAPC inferred three sub-populations (Figure 2d, Supplemental Figure S1ab). The minimal BIC values were obtained with *k* values ranging from 3 to 6, consistent with the optimal number of clusters determined by FastStructure. Comparing both methods, 95.3% and 90.0% concordant assignment were observed for *k* values of 3 and 6, respectively (Supplemental Figure S3).

Relatedness analysis among individuals revealed low intra- and inter-cluster genetic diversity, with accessions appearing neither significantly similar nor significantly distant (Supplemental Figure S4). This is consistent with the cumulative variance explaining genetic variation in the population, showing gradual increase with number of retained PCs up to 176 PCs (number of accessions in the GWAS panel) rather than reaching a plateau (Supplemental Figure S1a). Despite the overall genetic homogeneity, significative differences were observed between clusters for traits such as SM, HM, height, NC and ILI (Supplemental Figure S5). In particular, cluster K3 exhibited significant differences from the cluster K1 for these five traits, while the cluster K2 displayed intermediate trait values between K1 and k3. The clusters also differed in terms of size and genetic diversity, as indicated by their respective Tajima’s D values of 1.5, 2.2 and 0.8 for cluster K1 (n=32), K2 (n=57) and K3 (n=87), respectively. A positive Tajima’s D value suggests that these population underwent a recent bottleneck event, resulting in a depletion of low-frequency alleles. Based on Tajima’s D value, the K2 cluster exhibited greater genetic diversity compared to K1 and K3. Due to limited information on the pedigree of these accessions, no correlation was observed between population assignment and geographic or germplasm origins.

The limited genetic diversity observed in cultivated drug-type cannabis has historically been attributed to intensive clandestine breeding practices since the 1970s^2^. Despite the limited genetic diversity, cannabis exhibits a remarkable phenotypic variation that are highly desirable for breeding programs. Hence, it could be hypothesized that a portion of the observed phenotypic variations in cannabis may be attributed to transcriptional variations, along with potential contributions from epigenetic factors. Both in plants and animals, various factors, including copy number variations (CNVs)^60^, epigenetic elements^61^, and the insertion/deletion of transposable elements (TEs) in gene regulatory regions^62^, were found to affect phenotypic diversity, particularly traits important in domestication and breeding^63^. Therefore, an associated SNP may be in strong LD with either a candidate gene, where an allelic variant alters the phenotype, or with a regulatory region that either enhances or suppresses the expression of the phenotype^64^. The constrained availability of germplasm resources and low genetic diversity observed in cannabis pose significant limitations for breeding, which, in turn, hinder innovation and the long-term sustainability of the crop^6^. In contrast to other crops where wild-type or landrace varieties are promising genetic pools to enrich genetic diversity in breeding programs^65^, the situation in cannabis is more complex. Although hemp-type and drug-type cannabis diverged genetically long ago, they still share a considerable common pool of genetic variation, limiting the ability to mine rare alleles^66^. Given the growing demand for cannabis products, there is a critical necessity to pinpoint suitable genetic resources that can not only support production but also serve as a source of genetic diversity to help ongoing breeding efforts^6^.

### Identification of genomic regions controlling key agronomic and morphological traits

The GWAS analysis was performed using the SUPER and BLINK methods, with the incorporation of population structure (P) and cryptic relatedness (K*) as covariates to minimize the risk of false-positive associations. In total, 33 markers associated with the nine traits were identified, with 26 markers attributed to SUPER and 21 markers to BLINK (Figure 3, Table 1). Among these, 11 high confidence associations were commonly identified with both SUPER and BLINK. Four of these high-confidence SNPs (SNP_11, 13, 15, and 20) demonstrated significant phenotypic impact, with the proportion of phenotypic variance explained (PVE) ranging from 18% to 45%. Interestingly, several SNPs associated with different traits were located in close proximity to each other. For instance, SNP_6 and _17 were found within a region of approximately 58 kb on chromosome 1 (Chr01:86477139-86534795), while SNP_15, _21 and _31 were situated within a region of about 98 kb on chromosome 1 (Chr01:87397312-87494979). Most interestingly, all of these SNPs were located on an interval of ∼1 Mb and were associated with SD, SM, CD, ILI and height. The majority of markers identified have a modest influence on the phenotype (PVE < 10%), with approximately half of them exerting a negative impact. A negative impact on the phenotype, such as smaller size or shorter time to flowering or maturation, is not necessarily a negative factor for breeding. For example, the high confidence SNP_15 was estimated to have a negative impact of around 18% on height, which can be advantageous for maximizing indoor cultivation, where smaller plants are preferred.

**Figure 3.**
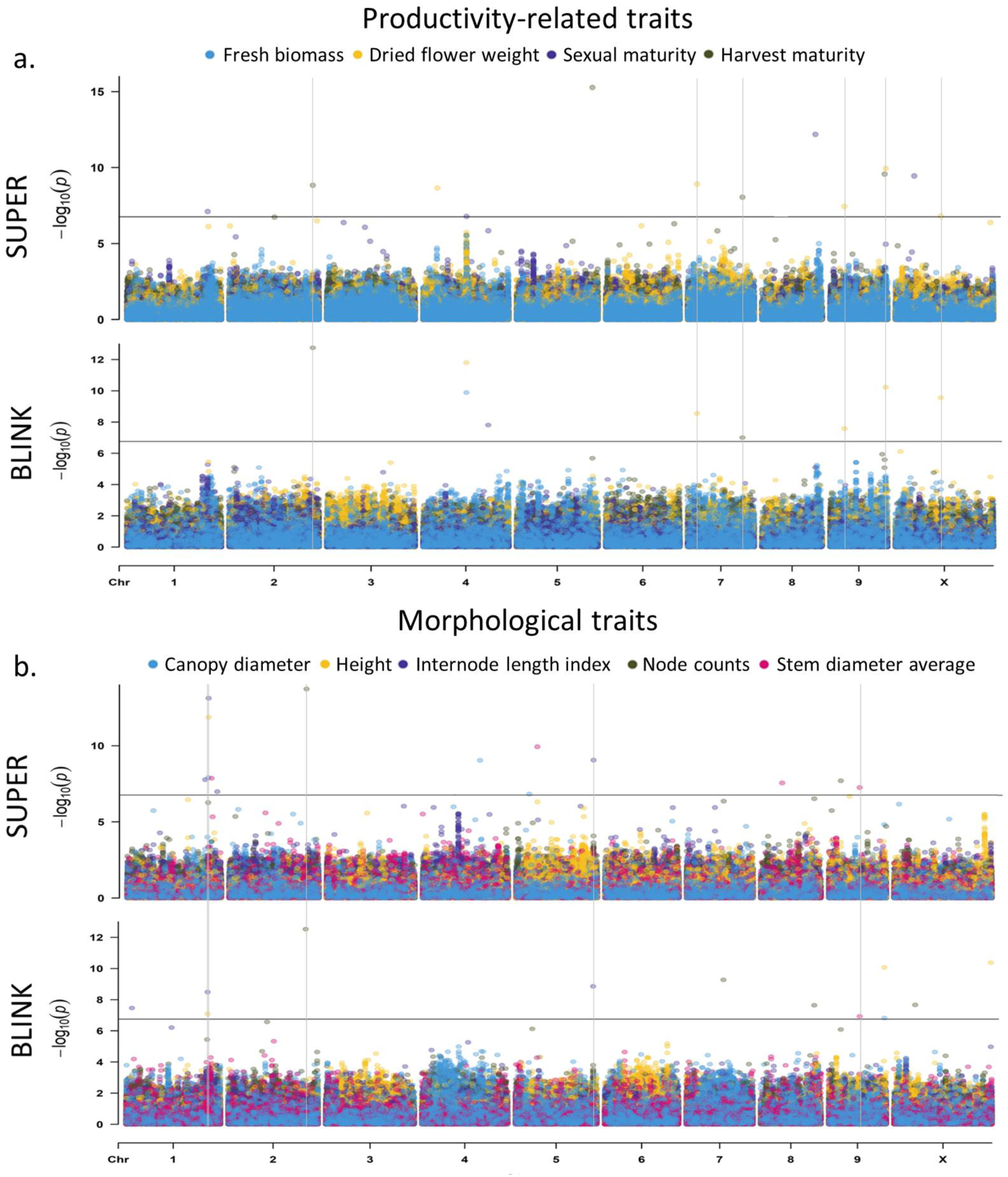
Genome-wide association studies (GWAS) for nine agronomic and morphological traits in drug-type cannabis. Manhattan plot for productivity-related traits **(a)** and morphological traits **(b)**. Each circle indicates the degree of association for a marker with a trait (y axis), while the x axis shows the physical position of each marker on a given chromosome across the genome. The horizontal grey line indicates the significance threshold (*p*-value=1.77 × 10^−7^, false discovery rate < 0.05). Settlement of MLM Under Progressively Exclusive Relationship (SUPER) and Bayesian-information and linkage-disequilibrium iteratively nested keyway (BLINK) are single- and multiple-locus analysis method.

**Table 1.** List of markers associated with nine different traits in drug-type cannabis identified through GWAS.

| Traits | Marker_ID | Chr. | MSS position | Major/minor allele | MAF<sup>a</sup> | *p*-value | PVE<sup>b</sup> | Effect | Model<sup>c</sup> |
| --- | --- | --- | --- | --- | --- | --- | --- | --- | --- |
| Fresh biomass | SNP_1 | 4 | 46549016 | G/A | 20.7% | 1.5E-10 | 27.2% | 107.77 | BLINK |
| Dried flower weight | SNP_1 | 4 | 46549016 | G/A | 20.7% | 1.6E-12 | 5.7% | 15.03 | BLINK |
|  | SNP_2 | 7 | 10007846 | A/G | 39.2% | 2.9E-09 | 3.0% | -10.18 | SUPER/BLINK |
|  | SNP_3 | 9 | 15308438 | C/T | 38.1% | 2.6E-08 | 5.4% | -10.39 | SUPER/BLINK |
|  | SNP_4 | 9 | 59690286 | T/A | 36.9% | 6.0E-11 | 4.9% | -13.34 | SUPER/BLINK |
|  | SNP_5 | X | 48657415 | C/T | 25.3% | 2.8E-10 | 8.4% | 15.11 | SUPER/BLINK |
| Sexual maturity | SNP_6 | 1 | 86534795 | T/G | 29.0% | 7.9E-08 | 6.4% | 3.84 | SUPER |
|  | SNP_7 | 4 | 46742383 | A/C | 10.5% | 1.7E-07 | 9.7% | 5.87 | SUPER |
|  | SNP_8 | 4 | 70020182 | C/A | 13.4% | 1.6E-08 | 4.3% | 7.02 | BLINK |
|  | SNP_9 | 8 | 57355911 | C/T | 32.1% | 6.5E-13 | 6.6% | -6.44 | SUPER |
|  | SNP_10 | X | 25733245 | C/T | 34.4% | 3.0E-08 | 3.4% | 2.29 | SUPER |
| Harvest maturity | SNP_11 | 2 | 89685715 | C/T | 8.8% | 1.8E-13 | 33.2% | 5.59 | SUPER/BLINK |
|  | SNP_12 | 5 | 81348921 | T/C | 8.0% | 5.3E-16 | 29.6% | -6.03 | SUPER |
|  | SNP_13 | 7 | 58735071 | C/T | 9.4% | 9.9E-08 | 35.3% | 4.20 | SUPER/BLINK |
|  | SNP_14 | 9 | 58422333 | C/T | 7.4% | 2.7E-10 | 18.9% | 4.68 | SUPER |
| Height | SNP_15 | 1 | 87494979 | T/C | 36.4% | 8.25E-08 | 17.97% | -3.67 | SUPER/BLINK |
|  | SNP_4 | 9 | 59690286 | T/A | 36.9% | 8.52E-11 | 6.14% | -3.32 | BLINK |
|  | SNP_16 | X | 103884160 | C/T | 31.8% | 4.18E-11 | 5.80% | 3.27 | BLINK |
| Stem diameter average | SNP_17 | 1 | 86477139 | T/C | 11.1% | 4.6E-09 | 3.8% | -0.02 | SUPER |
|  | SNP_18 | 5 | 22969509 | A/G | 9.9% | 1.2E-10 | 13.7% | 0.02 | SUPER |
|  | SNP_19 | 8 | 22916156 | G/A | 9.7% | 2.8E-08 | 3.1% | 0.02 | SUPER |
|  | SNP_20 | 9 | 33123470 | A/T | 7.1% | 1.2E-07 | 44.9% | 0.03 | SUPER/BLINK |
| Canopy diameter | SNP_21 | 1 | 87397312 | A/T | 40.1% | 1.24E-08 | 6.2% | 1.23 | SUPER |
|  | SNP_22 | 4 | 61731980 | A/C | 31.0% | 9.16E-10 | 4.8% | -1.75 | SUPER |
|  | SNP_23 | 5 | 14187264 | C/A | 27.0% | 1.50E-07 | 5.4% | 1.54 | SUPER |
|  | SNP_4 | 9 | 59690286 | T/A | 36.9% | 1.50E-07 | 24.3% | -1.93 | BLINK |
| Node counts | SNP_24 | 2 | 83198483 | G/C | 38.1% | 3.0E-13 | 7.6% | -2.32 | SUPER/BLINK |
|  | SNP_25 | 7 | 39816506 | C/- | 27.6% | 5.4E-10 | 6.3% | -1.67 | BLINK |
|  | SNP_26 | 8 | 57485105 | A/G | 32.4% | 2.3E-08 | 6.6% | -2.04 | BLINK |
|  | SNP_27 | 9 | 12818446 | A/C | 33.2% | 2.0E-08 | 11.3% | 1.81 | SUPER |
|  | SNP_28 | X | 22613259 | A/G | 14.8% | 2.2E-08 | 5.5% | -2.17 | BLINK |
| Internodes lenght index | SNP_29 | 1 | 6322667 | C/- | 29.0% | 3.4E-08 | 3.5% | 0.14 | BLINK |
|  | SNP_30 | 1 | 83867545 | C/A | 10.2% | 1.7E-08 | 3.7% | -0.21 | SUPER |
|  | SNP_31 | 1 | 87456694 | T/C | 35.5% | 3.3E-09 | 5.8% | 0.20 | SUPER/BLINK |
|  | SNP_32 | 1 | 96898415 | C/T | 18.8% | 1.0E-07 | 8.0% | -0.12 | SUPER |
|  | SNP_33 | 5 | 83258255 | A/G | 33.8% | 1.4E-09 | 5.8% | 0.19 | SUPER/BLINK |

Considering high-confidence SNPs does not preclude the relevance of other associated SNPs. For example, SNP_1 was only identified by BLINK but its impact on FB can be of particular interest for breeders compared to the high-confidence SNP_2 which has only a slight influence on DFW. Similarly, SNP_4 was high-confidence with DFW despite a modest effect on this trait. However, it notably reduces canopy size and slightly the height, which can help maximize plant density in cultivation. In both cases (i.e., high-confidence or not), as the identification of high-value markers for cannabis is in its early stages, the practical implementation of these markers by breeding programs will nevertheless require preliminary cross-validation. This can be achieved through meta-GWAS^67^, QTL mapping with biparental population and BSA. Additionally, comprehensive functional analyses of the candidate genes will be crucial. The utilization of both SUPER and BLINK in parallel maximizes the ability to detect a wide range of genetic associations. SUPER was used for single-locus GWAS, while BLINK can capture complex interactions between several loci through multi-locus analysis. This combined approach helps reduce false positives associations by allowing for cross-confirmation of results^51,52^. Furthermore, these models were each ranked as the most statistically powerful models for single and multi-locus analyzes for GWAS analyses in plants^20,50,52^. Overall, this approach revealed 33 associated markers, with one-third of them being high-confidence (identified by two models) and another one-third exerting interesting effects on the phenotype.

### Investigation of putative candidate genes

Among the 33 associated SNPs, 19 were in high LD (r²≥0.75) with other SNPs, forming 18 HBs (Table 2). Notably, SNP_15 and _31 were part of the same HB, spanning ∼97 kb on chromosome 1. The SNP_9, _17 and _26 were located within *LOC115699506*, *LOC115706481* and *LOC115699444*, respectively, without forming HBs. The 18 HBs spanned ∼420 kb, within which 43 annotated genes were identified. Consequently, these genes were considered as putative candidates genes associated with different traits. Recent genome annotation of cs10^44^ facilitated the investigation of the functions of candidate genes (Table 2). An orthology analysis was conducted by comparing the protein sequences of candidate genes with the Arabidopsis proteome^54^. Functional annotations were similar for the majority of candidate genes and their respective orthologs, confirming the robustness of the functional annotation of the cs10 transcriptome.

**Table 2:**
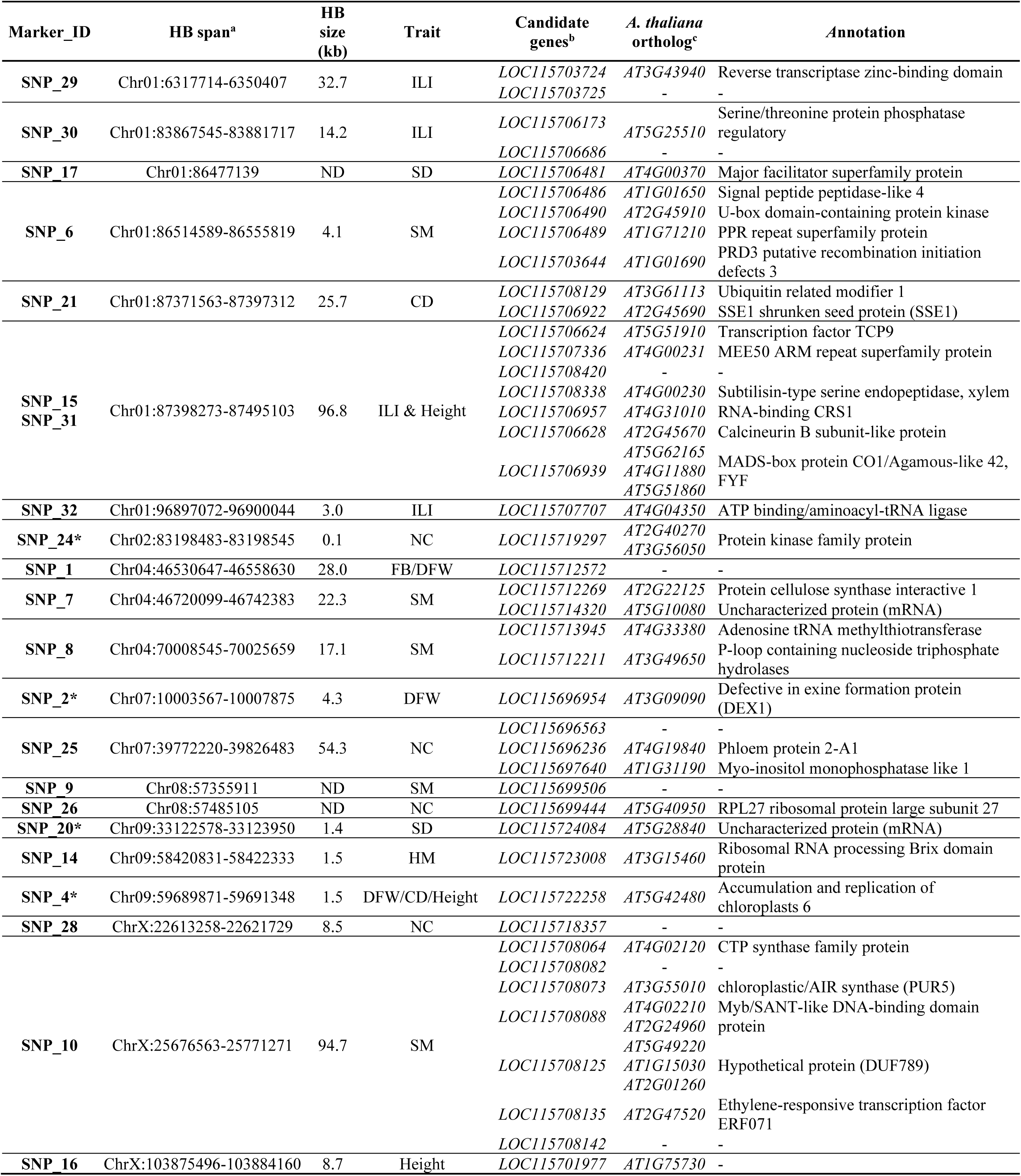
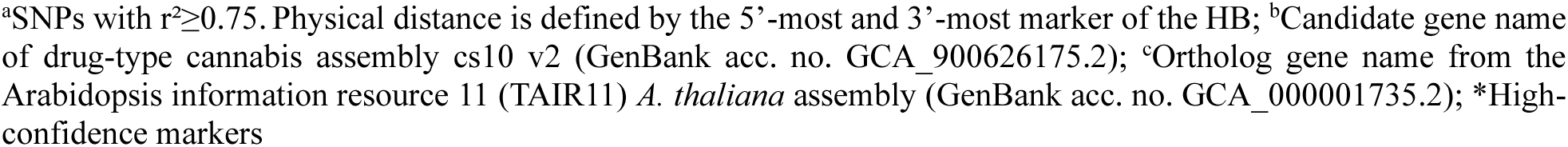
Orthology analysis of candidate genes located within haplotype block regions of markers associated with different traits.

The high-confidence SNP_4, which showed associations with DFW, CD, and height, was found to be in high LD with *LOC115722258*, associated with chloroplast metabolism and mechanisms. This suggests a potential link between the genetic variation of SNP_4 and the observed variations in these morphological traits through their impact on chloroplast-related processes. Furthermore, SNP_10, associated with DFW, was in high LD with *LOC115708088* and *LOC115708135*, annotated as having potential functions related to development and flowering^68,69^. In addition to structural genes, regulatory genes, such as transcription factors, were identified among the candidate genes (e.g., *LOC115706624* and *LOC115708135*). Approximately one-third of the associated SNPs were not in high LD with putative candidate gene, but they might more likely linked to gene regulatory regions. The *in-silico* identification of regulatory regions and their interaction with a gene is challenging and complex to link associated SNPs and the phenotype. However, this does not diminish their importance, especially for markers like high-confidence SNP_11 and _13, which were associated with a substantial impact on HM (PVE>30%). These findings suggest that regulatory elements, such as transcription factors, may play a critical role in shaping the phenotypic variation in cultivated cannabis.

## Conclusion

In conclusion, this study marks a pioneering exploration of the genetic landscape of Canadian drug-type cannabis through a comprehensive GWAS analysis, enriched by high-throughput genotyping and precise agronomic phenotyping data. Our findings open new avenues for advancing cannabis breeding programs and addressing the diverse needs of emerging industries. The application of a high-density genotyping approach yielded an extensive catalog of high-quality SNPs, effectively capturing the genomic diversity of drug-type cannabis. The well distribution of these markers across different chromosomes, coupled with high quality phenotypic data, facilitated the identification of molecular markers associated with complex agronomic and morphological traits. These markers hold great promise for further investigations to elucidate their functional links with phenotype variations, making them valuable assets for precision breeding efforts.

As we move forward, this research paves the way for in-depth studies to uncover the biological mechanisms governing these traits, potentially uncovering hidden genetic potential within cannabis populations. Furthermore, the implications of our work extend beyond immediate applications, as the identified markers are poised to play a pivotal role in the development of tailor-made cannabis cultivars, spanning both medicinal and recreational sectors, capable of meeting the dynamic demands of rapidly evolving industries.

Future perspectives in this domain encompass a deeper exploration of the candidate genes associated with the identified markers, seeking to unravel the intricate genetic and molecular underpinnings of these key traits. Additionally, functional validation experiments and expression profiling could elucidate the precise mechanisms through which these markers exert their effects. Collaborative efforts between academia and industry are essential to harness this newfound genetic knowledge and translate it into practical breeding strategies, ensuring the continued innovation and sustainability of the cannabis crop.

## Abbreviations

THC: Δ^9^-tetrahydrocannabinol
ICGRC: International Cannabis Research Consortium
NGS: Next-Generation Sequencing
GWAS: Genome-Wide Association Study
QTL: Quantitative Trait Loci
LD: Linkage Disequilibrium
RAD-Seq: Restriction-site Associated DNA Sequencing
GBS: Genotyping-By-Sequencing
MAS: Marker-Assisted Selection
GS: Genomic Selection
WGS: Whole-Genome Sequencing
HD-GBS: High-Density GBS
FB: Fresh Biomass
DFW: Dried Flower Weight
SM: Sexual Maturity
SD: Stem Diameter
CD: Canopy Diameter
ILI: Internode Length Index
NC: Node Counts
MAF: Minor Allele Frequency
HB: Haplotype Block
DAPC: Discriminant Analyses Of Principal Components
SUPER: Settlement of MLM Under Progressively Exclusive Relationship
BLINK: Bayesian-information and Linkage-Disequilibrium Iteratively Nested Keyway
ANOVA: Analyse of Variance
PERMANOVA: Permutational ANOVA
GO: Gene Ontology
QQ: Quantile-Quantile
PVE: Phenotypic Variance Explained

## Author Contributions

Maxime de Ronne: Conceptualization and Methodology; Data curation and analysis; Investigation; Visualization; Writing-original draft. Éliana Lapierre: Preparation of phenotypic data. Davoud Torkamaneh: Funding acquisition; Conceptualization and Methodology; Supervision; Writing-review. All authors have reviewed and approved the manuscript.

## Acknowledgments

The authors wish to thank Fuga Group Inc. for supporting this project. The authors also extend their sincere appreciation to Justine Richard-Giroux, Rosemarie Boulanger and Sean Kyne for their valuable contributions to the tedious phenotyping of the GWAS-panel.

## Ethics declarations

Not applicable.

## Data statement

The VCF files generated from the sequencing data and used for the analyzes of this study are on FigShare.com (and will be accessible after acceptance of the manuscript. This includes the raw SNP data set for the 176 accessions, the 282K imputed and filtered SNPs and the subdivision of the population by K clusters.

## Supplementary Material

**Supplemental Table S1.** Description of agronomic and morphological traits used in this study sourced from Lapierre et al. (2023b).

**Supplemental Table S2.** Description of filtering process and retained markers.

**Supplemental Table S3.** Distribution of markers across cannabis genome, LD and haplotype block (HB).

**Supplemental Figure S1.** Discriminant analysis of principal components (DAPC) of the 176 drug-type accessions using the whole 282K high-quality SNPs. a. Cumulated variance explained by the eigenvalues of the PCA. **b.** Value of BIC versus for increasing values of k. **c.** Optimization α-score graph. The optimal α-score suggested that twenty PCs should be retained for the discriminant analysis. **d.** DAPC cross-validation test for the optimal number of PCs retained.

**Supplemental Figure S2.** Population structure analysis of the 176 drug-type accessions using the whole 282K high-quality SNPs. Admixture plot for K values from 3 to 6 using FastStructure. The vertical lines represent the cannabis accessions, and the y-axis represents the probability that an individual belongs to a subgroup.

**Supplemental Figure S3.** Concordance of cluster assignment for FastStructure and DAPC inference for (a) *k*=3 and (b) *k*=6.

**Supplemental Figure S4. Heatmap of pairwise kinship matrix values between the 176 drug-type accessions.** The color histogram shows the distribution of co-ancestry coefficients. The redder the color, the more similar the individuals are, whereas the more yellow the color, the more dissimilar the individuals are.

**Supplemental Figure S5.** Distribution of agronomic traits for 176 drug-type accessions by K assignments. Letters represent significant difference according to Tukey’s HSD test.

**Supplemental Figure S6.** Quantile-quantile (QQ) plots of genome-wide association analysis with different statistical methods and traits.

